# Intron-driven gene expression in the absence of a core promoter in *Arabidopsis thaliana*

**DOI:** 10.1101/083261

**Authors:** Jenna E Gallegos, Alan B Rose

## Abstract

In diverse eukaryotes, certain introns increase mRNA accumulation through the poorly understood mechanism of intron-mediated enhancement (IME). A distinguishing feature of IME is that these introns have no effect from upstream or more than 1 Kb downstream of the transcription start site (TSS). To more precisely define the intron position requirements for IME in Arabidopsis, we tested the effect of the *UBQ10* intron on gene expression from 6 different positions surrounding the TSS of a *TRP1:GUS* fusion. The intron strongly increased expression from all transcribed positions, but had no effect when 204 nt or more upstream of the 5’-most TSS. When the intron was located in the 5’ UTR, the TSS unexpectedly changed, resulting in longer transcripts. Remarkably, deleting 303 nt of the core promoter, including all known TSS’s and all but 18 nt of the 5’ UTR, had virtually no effect on the level of gene expression as long as a stimulating intron was included in the gene. When the core promoter was deleted, transcription initiated in normally untranscribed sequences the same distance upstream of the intron as when the promoter was intact. Together, these results suggest that certain introns play unexpectedly large roles in directing transcription initiation and represent a previously unrecognized type of downstream regulatory elements for genes transcribed by RNA polymerase II. This study also demonstrates considerable flexibility in the sequences surrounding the TSS, indicating that the TSS is not determined by promoter sequences alone. These findings are relevant in practical applications where introns are used to increase gene expression and contribute to our general understanding of gene structure and regulation in eukaryotes.

## Introduction

The transcription start site (TSS) of genes transcribed by RNA Polymerase II in eukaryotes is thought to be primarily determined by the assembly of the pre-initiation complex on recognizable sequences in the core promoter (reviewed in (1, 2, 3)). This process starts with binding of the TATA-binding protein subunit of the general transcription factor TFIID to the TATA box sequences 30-40 nt upstream of the TSS. Core promoters often contain additional recognizable sequences including the initiator, which encompasses the TSS, the TFIIB recognition element, and the downstream promoter element, which, along with the initiator, binds to TFIID (4, 1). However, there are no universally conserved core promoter elements, and even the TATA box is found in a minority of genes in plants (5), yeast (6), and humans (7). The common heterogeneity in start sites within a single gene further suggests that there is considerable flexibility in the sequences that can support transcription initiation (5). In addition to the core promoter, many genes rely on proximal promoter elements (usually less than 1 Kb from the TSS) and distal enhancer elements to regulate gene expression (3).

There are many examples where a fully intact promoter with all its transcription factor binding sites is inactive unless one or more of the gene’s endogenous introns are included (8, 9, 10, 11, 12). The expression of other genes that are weakly active without an intron can be increased tenfold or more by the addition of an intron (8, 13, 14, 15, 16). Furthermore, introns can change the spatial expression patterns of tissue-specific genes (12, 17, 18). This demonstrates that some introns significantly boost expression, even in the absence of prior promoter activity.

In some cases, the mechanism through which an intron increases gene expression is well understood. For example, introns can contain enhancers or alternative promoters (19, 20, 21). In addition, interactions between the splicing machinery and other factors involved in mRNA synthesis and maturation, as well as the exon-junction complex proteins deposited on the mRNA during splicing, assist in mRNA production, stability, export, and translation (22, 23, 24, 25).

Certain introns that stimulate gene expression exhibit properties that suggest that they must operate by a different and poorly understood mechanism. First, the varying abilities of efficiently spliced introns to increase mRNA accumulation indicate that only certain introns boost mRNA levels beyond the general effects caused by the splicing machinery or exon junction complexes (14). Second, these introns are unlike enhancers because they must be located within 1kb of the TSS to have an effect (26). Third, the sequences responsible for increasing mRNA accumulation are redundant and dispersed throughout these stimulating introns (27), unlike the discrete sequences to which transcription factors bind in promoters and enhancers. The increase in mRNA accumulation caused by specific introns only when in transcribed sequences near the TSS will be referred to here as intron-mediated enhancement (IME) (28).

An algorithm known as the IMEter assigns a score based on the oligomer composition of a given intron compared to promoter-proximal introns genome-wide (23). The utility of the IMEter in predicting the stimulating ability of introns has been confirmed in Arabidopsis (27), rice (29), soybeans (30), and other angiosperms (31). The sequence TTNGATYTG is over-represented in introns with high IMEter scores (27) and is sufficient to convert the previously non-stimulating *COR15a* intron into one that strongly boosts mRNA accumulation. Introns containing either 6 or 11 copies of this motif, named *COR15a6L* and *COR15a11L*, increase mRNA accumulation 15-fold and 24-fold respectively (32).

While the ability of an intron to stimulate expression is known to vary with its location within a gene, the exact positional requirements for IME have not been fully determined. Early studies demonstrated that most introns that boost expression from the 5’ end of a gene have no effect when inserted into the 3’ UTR (8, 17, 33). Experiments varying the location of an intron in *TRP1:GUS* coding sequences revealed that their ability to stimulate mRNA accumulation declines when moved from roughly 250 nt to 550 nt downstream of the major TSS, and is completely gone when 1100 nt or more form the major TSS (26). In most previous studies in which the intron was placed upstream of the promoter (8, 34, 14), the distance between the intron and the TSS was more than 550 nt, so the introns may have been too far away to affect expression. While these experiments clearly demonstrated that introns are unlike enhancer elements whose influence extends several kilobases, they did not establish if introns must be transcribed to have an effect, or if they must simply be near the TSS to boost expression whether transcribed or not.

Furthermore, it is unknown if the stimulating ability of an intron continues to increase as it gets closer to the TSS, or if introns have their maximum effect approximately 200 nt downstream of the major TSS where genome-wide average intron IMEter scores peak (35). Pinpointing the ideal location for a stimulating intron will help to maximize gene expression in industrial and research applications. Additionally, the differing effects of a stimulating intron at various locations could yield insight into the mechanism of IME.

In this study, we completed the first gene-scale mapping of the effect of intron location on IME by comparing the expression of *TRP1:GUS* fusions containing the *UBQ10* intron at different locations around the transcription start site. In the process, we discovered an unexpected role for introns in determining the site of transcription initiation.

## Results

### The 5’-limit of intron position for IME

To determine the 5’-limit from which an intron can stimulate expression by IME, the first intron of the *UBQ10* gene was tested at six locations upstream of the normal second exon of a *TRP1:GUS* reporter gene (designated positions 1-6 in Figure 1a and Supplementary Figure 1). Previously generated (26) transgenic *TRP1:GUS* plants with the first intron of *UBQ10* at three additional locations, designated position 0, −1, and −2, were included in figure 1 to show the 3’ limits of intron position for IME. Position 0, −1, and −2 had been previously designated “259”, “551”, and “1136” respectively according to their distance from the most frequently used transcription start site. A different numbering system was used here because a description of intron location relative to the TSS is complicated by the presence of multiple TSSs in this gene. The main transcription start site is at −41 relative to the ATG, as determined by primer extension and RNAse protection (36), RNAseq, and the 5’ ends of ESTs and cDNAs in the TAIR database (https://www.arabidopsis.org/). There are also a few minor transcription start sites, of which the one at −117 is the furthest from the start codon and is the annotated TSS for this gene.

**Figure 1.**
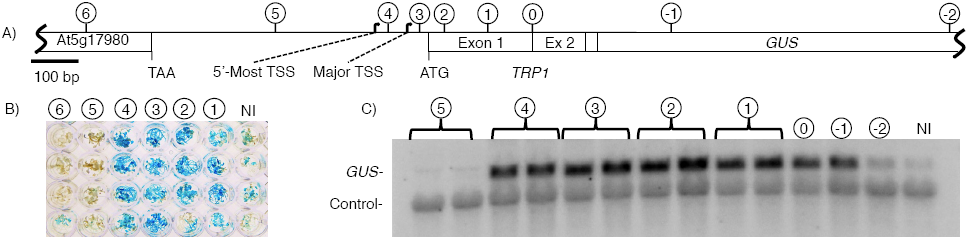
Comparing the stimulating ability of the *UBQ10* intron at locations near the TSS. **A)** The first intron of *UBQ10* was inserted at one of six locations (numbered 1-6) around the TSS. Previously generated constructs with the same intron at position 0, −1, and −2 were also included to show the full limits of intron position for IME. Arrows mark the most commonly used TSS (-41 relative to the start codon), and the 5’-most TSS (-117 relative to the start codon). **B)** Histochemical staining for *GUS* activity in transgenic plants that contain the *TRP1:GUS* fusion with the *UBQ10* intron at the indicated position or no intron (NI). Each well in a vertical column contains five T_2_ seedlings from an independent line of unknown copy number. **C)** RNA gel blot probed with *GUS* and a loading control (the endogenous *TRP1* gene). Each adjacent lane with the same label represents an independent single-copy homozygous line.

Introns inserted at position 3 should be in the 5’-UTR of all *TRP1:GUS* transcripts, while position 4 is upstream of the major TSS but downstream of the 5’-most TSS. The intron at position 5 is upstream of all known TSSs, and roughly midway between the *TRP1* start codon and the stop codon of the upstream gene. This gene (*At5g17980*) is a 3.15 Kb intronless gene of which only 1.8 Kb from the 3’ end is present in the *TRP1* promoter fragment in the *TRP1:GUS* fusions. The intron at position 6 is in coding sequences of this gene 198 nt from the stop codon.

To insert the intron, a *PstI* site was added to the 5’ end of the intron, and the last six nucleotides of the intron were converted into a *PstI* site. The intron was then cloned as a *PstI* fragment into a *PstI* site created by site directed mutagenesis at the desired location (Supplemental Figure 1) as described (37). Introns inserted as *PstI* sites leave no extraneous nucleotides in the mRNA (26), and are efficiently spliced, presumably because the 5’ splice site is unchanged, and the 3’ *PstI* site (CTGCAG) conforms to the major 3’ splice site consensus (NNNYAG) in dicots.

For intron positions in coding sequences (positions 1 and 2) there were no places where *PstI* sites could be made without changing an amino acid, so sites were chosen such that changes were unlikely to disrupt function. The *PstI* sites introduced at positions 1 and 2 convert an isoleucine and a glycine respectively into alanines, all of which are in the family of nonpolar amino acids. Further, the first two exons of the *TRP1* gene encode the chloroplast transit peptide (36). Transit peptides have non-stringent sequence requirements and can usually tolerate moderate changes without losing function (38). They are also cleaved off as the protein is imported into the chloroplast, so conservative changes here are unlikely to affect the enzymatic activity of the *GUS* reporter (37).

To qualitatively assess the positional effect of the *UBQ10* intron on expression of the *TRP1:GUS* reporter, pooled T_2_ transgenic seedlings of unknown transgene copy number were histochemically stained for GUS activity with X-gluc (Figure 1b). The *UBQ10* intron clearly stimulated expression from all four transcribed locations but not from either position upstream of the TSS where they may reduce expression. This is consistent with previous observations that introns acting by IME fail to increase expression from upstream of the promoter (8), and suggests that introns may need to be transcribed to affect expression, even if they are within a few hundred nucleotides of the TSS.

To quantitatively compare expression of reporter genes containing the intron at all transcribed positions, GUS enzyme activity and mRNA accumulation were measured in single-copy transgenic *A. thaliana* lines (Figure 1c, and Supplementary Tables 1 and 2). Expression levels between independent single-copy lines of each construct were similar indicating that, for this transgene, the site of integration had little effect. This has been demonstrated for other *TRP1:GUS* fusions (37, 26), and likely stems from the fact that the *TRP1* promoter is in the middle of the T-DNA insertion and is isolated from flanking plant sequences by at least 2 Kb of sequence on each side.

*GUS* enzyme assays and RNA gel blots both indicated that *TRP1:GUS* mRNA levels were similar when the *UBQ10* intron was located at position 0, 1, 2, 3, or 4 (Figure 1, and Supplementary Tables 1 and 2). The activity of the intron at position 4 was surprising because this location is 43 nt upstream of the major TSS, meaning that most of the transcripts were not predicted to contain the intron. Inserting 304 nt of sequence between the core promoter and the major TSS was expected to reduce expression.

### The effect of intron position on TSS location

To determine if any transcripts initiated within the intron, cDNA from pooled seedlings was amplified by 5’-RACE, cloned, and sequenced. None of the sequenced products contained any intron sequences, but they did reveal an effect on the site of initiation. When the intron was located at positions 1 or 2 (within the first exon), the 5’ ends of most 5’-RACE products, and presumably the TSS, mapped to within 10 nt of the major TSS (Figure 2). However, when the intron was located at position 3 (within the 5’-UTR), transcription began almost exclusively within 5 nt of the furthest upstream TSS (Figure 2). When the intron was located at position 4 (upstream of the major TSS), transcription began at the 5’-most TSS or at locations further upstream where initiation does not normally occur, but not at any site downstream of the intron including the major TSS (Figure 2). This suggests that introns may affect the site of transcription initiation and explains how the gene could still be highly expressed when the intron was located upstream of the major TSS.

**Figure 2.**
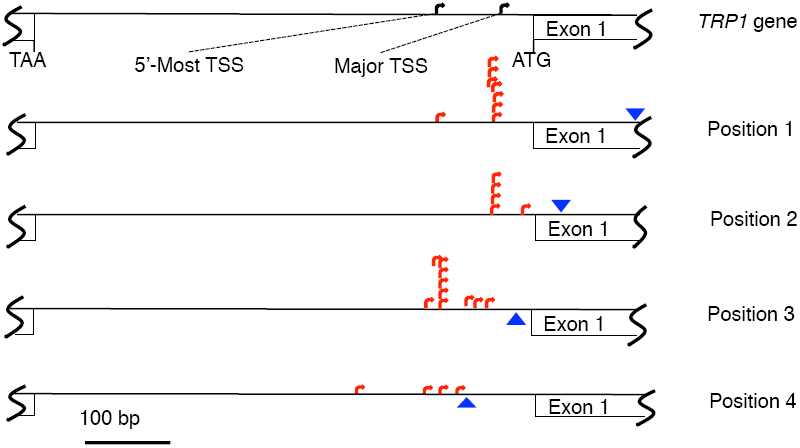
Locations of the 5’-end of *TRP1:GUS* 5’-RACE products when the *UBQ10* intron was at position 1, 2, 3, or 4 (intron position marked with blue triangles). Each red arrow represents an individual sequenced cDNA.

### Expression in the absence of the core promoter

The observation that inserting the intron into the 5’-UTR changed the location at which transcription predominantly began suggests that transcription does not always start a fixed distance downstream of conserved sequences in the core promoter. To determine the extent of the flexibility in sequences that can support initiation, 303 nt of the *TRP1* core promoter were deleted from the *TRP1:GUS* fusion (from position 3 to 5 in Figure 1a). The deletion encompassed all previously recognized transcription start sites and all but 18 nt of the 5’-UTR (Supplementary Figure 1). The deletion was created in reporter gene fusions containing no intron, the non-stimulating *COR15a* intron, or one of three stimulating introns: *UBQ10*, *COR15a6L*, or *COR15a11L*. Remarkably, deleting the core promoter had no obvious effect on gene expression, as determined by histochemical staining for *GUS* activity in seedlings of unknown transgene copy number (Figure 3a). To quantify potential subtle differences in expression, mRNA levels were compared in single-copy transgenic *A. thaliana* lines. Deleting the promoter did not diminish mRNA levels from *TRP1:GUS* fusions containing any of the three stimulating introns (Figure 3b). However, deleting the promoter reduced but did not eliminate expression of the constructs containing the non-stimulating *COR15a* intron or no intron (Figure 3b). The size of the mature mRNA was not changed by the promoter deletion regardless of which intron was present.

**Figure 3.**
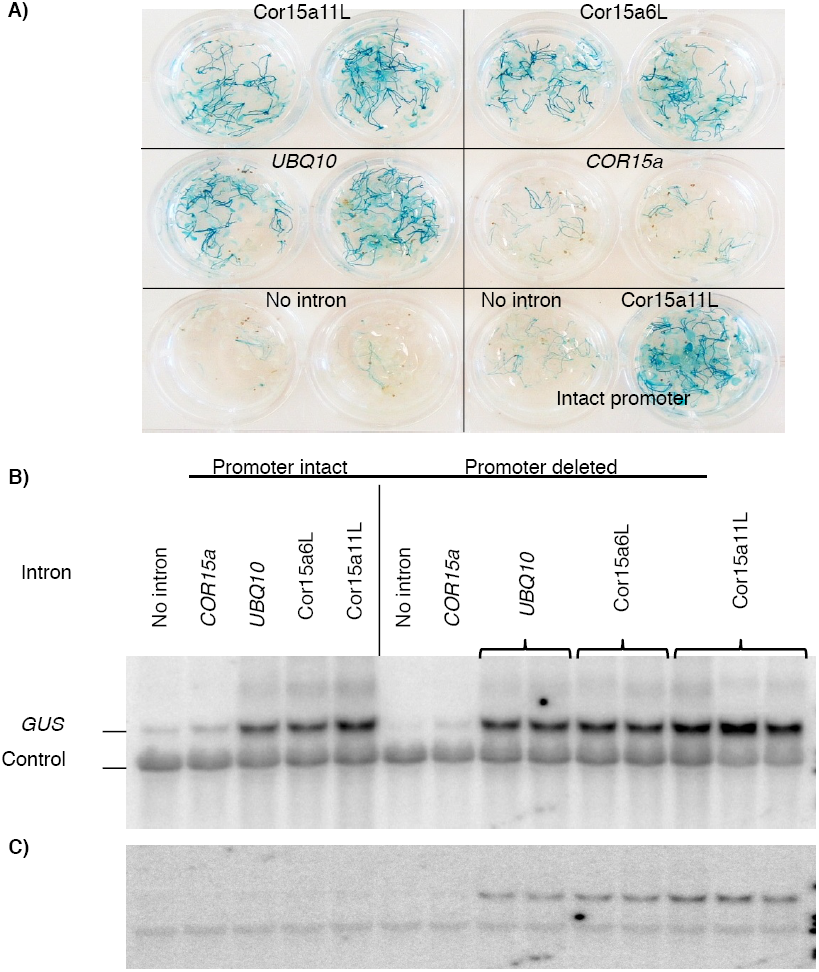
Introns stimulate expression in the absence of the core promoter. **A)** Histochemical staining of transgenic plants containing *TRPI:GUS* fusions with the indicated introns. The promoters in all constructs except the two on the bottom right have had all sequences between positions 5 and 3 deleted. Each circular well contains 20 unrelated T_1_ seedlings from an independent transformation. **B)** RNA gel blot probed with *GUS* and a loading control (the endogenous *TRPI* gene). Each lane with the same label contains RNA from an independent single-copy homozygous T_3_ line. **C)** Same blot as in **B)** probed with a *TRP1* promoter fragment extending from positions 4 to 6.

### TSS location in the absence of the core promoter

To determine where transcription was initiating in the deletion-containing constructs, and to verify that transcription of the *TRP1:GUS* fusion was similar to that of the endogenous *TRP1* gene, cDNA from pooled seedlings was amplified by 5’-RACE, cloned, and sequenced. For *TRP1:GUS* constructs containing the *COR15a11L* intron in which the promoter was intact, the apparent transcription start sites mapped within the range of −114 to +1 relative to the ATG (Figure 4), consistent with the start sites of the endogenous *TRP1* gene.

**Figure 4.**
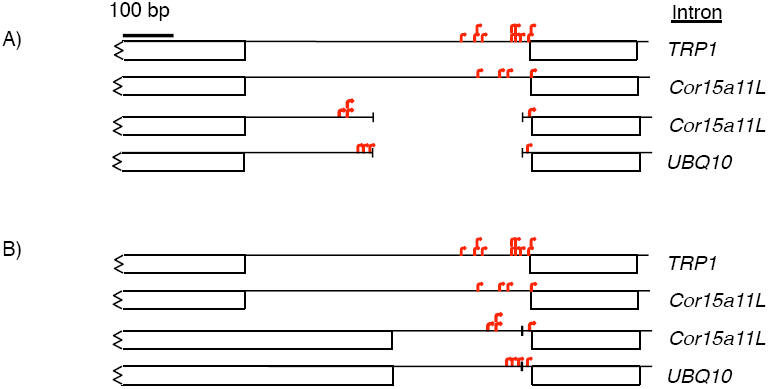
Deleting the core promoter changes the TSS. **A)** Locations of TSSs of the *TRP1* gene (top line) and *TRP1:GUS* fusions containing the indicated introns, with the promoter intact (second line) or deleted (bottom two lines). The sequences are aligned to the genome, so the promoter deletion appears as a gap. The *TRP1* TSSs were determined by primer extension and RNAse protection (36) and from cDNAs and ESTs in the TAIR database (https://www.arabidopsis.org/). The *TRP1:GUS* TSSs were determined by 5’-RACE, with each red arrow representing an individual sequenced cDNA. **B)** As in **A)**, but with the gap closed to show the distances of the TSS’s from the start of the intron.

When the promoter was deleted, most of the apparent start sites from constructs containing either the *COR15a11L* or the *UBQ10* intron mapped upstream of the deletion in normally untranscribed sequences (Figure 4). Even though these initiation sites are far upstream of the normal TSS, they are a similar distance upstream of the intron (223-311 nt) as when the promoter was intact. This can be seen in the average length of the 5’-UTRs in the sequenced 5’-RACE clones, which did not significantly differ between the promoter-deletion constructs containing either the *Cor15a11L* or *UBQ10* intron, the intact promoter with the *Cor15a11L* intron, or the previously sequenced cDNAs and ESTs from the endogenous *TRP1* gene.

To verify that deleting the core promoter caused transcription to start in a new location, RNA gel blots were probed with a 709 nt fragment spanning between the location of *PstI* sites at positions 4 and 6 (Figure 3c). The probe hybridized much more strongly with mRNA from lines in which the promoter was deleted and a stimulating intron was present than with the promoter intact regardless of whether or not an intron was present. This confirms that transcripts derived from promoter deletion constructs with stimulating introns contain upstream sequences that are not transcribed when the promoter is intact. The smaller band present in all lanes is of unknown origin.

### Expression in the absence of the entire intergenic region

To further test the limits of sequences that can support transcript initiation, a larger deletion (spanning from position 3 to position 6 in Figure 1a) was created in *TRP1:GUS* fusions containing no intron or the *UBQ10* intron. This deletion removed the entire intergenic region from 18 nt upstream of the *TRP1* ATG into coding sequences near the 3’-end of the upstream gene, and encompassed the whole region of DNAse sensitivity associated with the endogenous *TRP1* promoter (Supplementary Figure 2). Transgenic plants containing *TRP1:GUS* constructs with this complete promoter deletion had undetectably low levels of *GUS* activity even when a stimulating intron was included (Supplementary Figure 3). This suggests that even though the promoter sequences that can support initiation in response to a stimulating intron are surprisingly flexible, there are some features of the *TRP1* promoter, either sequences or chromatin structure, that are absolutely required for expression.

## Discussion

### Intron position and gene expression

With the results presented here, the effect on expression of the *UBQ10* intron has been measured from a total of 14 locations within the *TRP1:GUS* reporter gene. The intron increased mRNA accumulation from only six positions near the start of the gene. The 5’- and 3’-most positions from which the intron boosted expression are 594 nt apart. Of the six active positions, the effect of the intron on mRNA accumulation was greatest at −18 from the start codon and least from +510, but the effect at positions 0-4 differed from the average by less than 25%. The intron position for maximum IME was closer to the start of transcription than predicted from average IMEter scores, which peak several hundred nucleotides downstream of the TSS. However, this could be due in part to TSS annotation errors in the genomic data used to analyze IMEter score distributions. In practical applications where introns are used to maximize gene expression, for both efficacy and ease it might be best to insert the intron into the 5’-UTR near the start codon.

It was previously unclear whether introns could affect gene expression if they were upstream of but near the TSS. Here we showed that the *UBQ10* intron, which strongly affects expression from 627 nt downstream of the 5’-most TSS (position 1), clearly did not stimulate expression when located 204 nt upstream of the 5’-most TSS (position 5).

While this finding is consistent with the idea that introns must be transcribed to have an effect, this conclusion is complicated by the possibility that stimulating introns might cause initiation upstream of themselves, as discussed below.

### Intron position and the TSS

The lack of universal promoter elements (2) illustrates a high degree of flexibility in the sequences that can support transcription initiation. Nonetheless, transcription tends to initiate within the same region for most genes. In genes with many TSSs, it remains unclear whether each TSS lies a fixed distance from multiple functional promoters, or if transcription starts at varying distances from a single promoter because polymerase scans a flexible distance before initiating. Our finding that moving a stimulating intron into the 5’-UTR, or deleting the core promoter, causes transcription to begin in sequences that do not normally support initiation suggests that start sites are not determined by promoter sequence alone. There must be additional conditions that can be influenced by an intron to allow initiation at competent sites that are normally inactive.

The transcription stimulated by the *UBQ10* intron did not initiate a fixed distance upstream of the intron, as moving the intron through coding sequences towards the start of the gene did not alter the TSS until the intron reached the 5’-UTR. Relocating the intron over a range of nearly 500 nt (at positions +29, +118, +218, and +510 relative to the ATG) in coding sequences of constructs with intact promoters did not appreciably change the TSS, as determined by the size of transcripts on RNA gel blots and sequencing 5’-RACE products (Figures 1, (2, and (26)). When at position 4, the intron separated the promoter from the main TSS, possibly leading to the preferential use of TSSs upstream of the intron at −117 and other locations. However, the predominant TSS also shifted when the intron was at position 3, downstream of the main TSS. In yeast, splicing occurs during transcription very soon after the nascent RNA emerges from RNA polymerase II (41), so moving the intron into the 5’-UTR might push the predominant site of initiation further upstream if there is steric interference between the bulky spliceosome and the transcription machinery as it assembles on the promoter for the next round of transcription.

### Promoter deletions

Deletions in the *TRP1* core promoter resulted in comparable levels of transcript initiation 220-310 nt upstream of the intron, as when the promoter was intact. The sequences between positions 5 and 3 are clearly dispensable for expression, even though this region contains all known TSSs for this gene. The larger deletion extending to position 6 in the upstream gene eliminated all expression regardless of the presence of introns. This indicates that the sequences between positions 5 and 6 are capable of supporting initiation, while those upstream of position 6 are not.

The region between positions 5 and 6 may have distinguishing features beyond specific sequence composition, such as a propensity to form open chromatin. The sequences between positions 3 and 6 are noticeably more AT-rich than sequences upstream of position 6, and histone associations are favored by GC-richness (42). Further, the DNAse sensitive region that includes the *TRP1* promoter does not extend as far upstream as position 6.

These results may help to explain why some promoters are completely dependent on introns for expression. A subset of intron-dependent genes may not have core promoters in the traditional sense but rather rely on introns to initiate transcription upstream of themselves. The ability of stimulating introns to override the tissue specificity of promoters further suggests that the mechanism of IME does not depend on prior promoter activity, and inherently leads to constitutive expression throughout the plant.

If introns can drive expression in the absence of a core promoter, it is puzzling that promoters have not been found more generally dispensable in the large number of publications reporting promoter deletion analyses. One possible explanation is that introns are rarely included in the genes used to assess promoter activity. One exception is a study of the *unc-54* gene of *Caenorhabditis elegans* in which the expression of different versions of the gene was measured by their ability to rescue an *unc-54* null mutation (43). While *unc-54* genomic DNA rescued efficiently, *unc-54* cDNA under the control of the *unc-54* promoter did not. An intron-containing but promoterless gene rescued, and transcripts derived from this construct initiated in the plasmid sequences newly fused upstream of the start codon. Either some bacterial sequences can fortuitously act as a promoter in *C. elegans*, or the first intron of *unc-54* causes initiation upstream of itself.

There is suggestive evidence from the ENCODE project that some introns might cause initiation upstream of themselves in humans as well. Using unique EST ends as a genome-wide indicator of transcription start sites, initiation was observed not only at the annotated transcription start site but also 250-300 nt upstream of the first intron of genes with long first exons (44). Additionally, genes with a naturally occurring intron very near the transcription start site tend to be more actively transcribed, as demonstrated by association with RNA Polymerase II and TFIID (44). Thus, the apparent ability of introns to stimulate transcript initiation upstream of themselves and for promoter proximal introns to increase gene expression may be conserved across kingdoms.

### Model

While the mechanism through which introns increase gene expression remains unclear, the results presented here indicate that introns can have an unexpectedly large influence on determining the location at which transcription initiates. Any model of IME must account for the locations from which introns affect expression, and the ability of some but not all upstream sequences to substitute for the normal transcription start sites of the *TRP1* gene. The following speculation incorporates the results presented here with previous data regarding the effect of introns on gene expression in plants and other eukaryotes. This discussion focuses largely on the *UBQ10* intron because it is the Arabidopsis intron for which the most data are available. From the number of introns with high IMEter scores, we predict the expression of as many as 15% of genes in Arabidopsis may be influenced by a similar mechanism (28). There are likely other ways in which different introns increase expression.

The main requirement for sequences to act as the promoter of intron-regulated genes such as *TRP1* might be a region of open chromatin that allows access of the transcription machinery to the DNA. Within that region of open chromatin there may be DNA sequences that favor initiation, but others can readily substitute when the preferred sites are deleted. Introns may also help to create the local chromatin state necessary for initiation through the interaction between DNA sequences within the intron and histone modifying or nucleosome positioning factors. Transcription from many or most promoters may be inherently bidirectional (45). In yeast, introns have been shown to affect the proportion of transcripts heading towards coding sequences by recruiting termination factors that decrease transcription in the opposite direction (46). Introns also favor re-initiation by facilitating DNA looping that brings the 3’ end of the gene, and thus the transcription machinery, in proximity to the promoter (47). In short, stimulating introns may boost mRNA production by creating local chromatin conditions that favor initiation, decreasing the proportion of PolII molecules that transcribe away from the gene, and positive feedback that amplifies the accumulation of mRNA through reinitiation.

The limitations to the positions from which the *UBQ10* intron is capable of stimulating mRNA accumulation could be caused by a combination of the relatively short distances (1 Kb or less) over which the postulated intron-driven changes in chromatin structure extend, and the location of potential translational start codons (as described (28)). Ribosomes usually scan from the cap structure at the 5’ end of an mRNA and initiate translation at the first ATG, and mRNAs that contain a termination codon too far from their 3’ end are rapidly degraded by nonsense-mediated mRNA decay (48, 49, 50). Transcripts that initiate either upstream or downstream of the 220 nt window in which the *TRP1* start codon is the first ATG are therefore likely to be unstable (Supplementary Figure 4). The intron at all locations might increase transcription initiation upstream of itself, but mRNA accumulates only when the intron is near the 5’ end of the gene because only then does the *TRP1:GUS* open reading frame occupy almost the entire mRNA. This might also explain why there was actually a slight decrease in expression when the intron was located at position 5 or 6 if transcription was initiating preferentially upstream of the intron. Both the deletion between sites 5 and 3 that permits expression, and the deletion between sites 6 and 3 that eliminates it, leave windows (112 nt and 187 nt respectively) in which transcription could initiate to generate functional TRP1: *GUS* protein. The apparent lack of change in TSS when the *UBQ10* intron is moved through *TRP1:GUS* coding sequences is consistent with the idea that only transcripts that start within the 220 nt window upstream of the ATG accumulate, and that within that range the preferred site for initiation is at −41.

### Conclusion

The ability of some introns to increase mRNA accumulation from more than 500 nt downstream of the TSS, but not from more than 1100 nt away, and to stimulate expression of a gene lacking its core promoter, indicates that introns represent a previously unrecognized type of regulatory element for genes transcribed by RNA polymerase II. The ability of an intron to effect gene expression over a range of several hundred nucleotides suggests that introns are unlike the downstream transcription factor binding sites found in genes transcribed by RNA Polymerase III. The flexibility in sequences that can support transcript initiation in response to an intron may contribute to the difficulty of identifying conserved promoter elements. Further characterization of the sequences that support initiation in response to an intron, and the role of chromatin structure, should lead to better understanding of the mechanism through which introns increase gene expression.

## Materials and Methods

### Cloning of reporter gene fusions

The starting template for all constructs was an intronless *TRP1:GUS* fusion containing 1.8 Kb of sequence stretching from the 3’ end of the gene upstream of *TRP1* (*At5g17980*) through the first 8 amino acids of the 3rd exon of *TRP1* fused to the *E. coli uidA* (*GUS*) gene (39). *PstI* sites were introduced at 6 locations using PCR mutagenesis, and confirmed by sequencing (26). For intron position experiments, the first intron of the *UBQ10* gene was inserted as a *PstI* fragment at each of the 6 locations (26). To delete the core promoter, a promoter fragment whose 3’ end was the *PstI* site at position 5 (in the intergenic region) was ligated to an exon fragment whose 5’ end was the *PstI* site at position 3 in the 5’-UTR, thereby deleting the sequences between positions 5 and 3. The same procedure was used to generate the full promoter deletion, except that the distal fragment had the *PstI* site at position 6 (in the upstream gene). All fusions were then cloned into the binary vector pEND4K, transformed into *Agrobacterium tumefaciens* by electroporation, and then introduced into Arabidopsis thaliana ecotype Columbia (Col) by floral dip as described (37).

### Qualitative *GUS* expression assays

For qualitative comparisons of expression (as in Figure 1b and Figure 3a), T_2_ seedlings for multiple lines for each construct were histochemically stained for *GUS* activity. The plate of seedlings in buffer containing the substrate 5-bromo-4-chloro-3-indolyl ß-D-glucuronic acid (Calbiochem, La Jolla, CA, USA) was incubated at 37° for 30 minutes to 2 hours, the plants were rinsed in water, and chlorophyll was removed with ethanol.

### Identification of single-copy transgenic lines

Single-copy transgenic lines were identified for quantitative measurements of *GUS* enzyme activity and mRNA levels. For each construct, 18-72 T_2_ lines were screened for segregation ratios by kanamycin resistance (26). Genomic DNA was extracted from lines that exhibited a 3:1 segregation ratio (kanamycin resistant:sensitive) indicative of a single locus of transgene insertion. The DNA was digested with *BamHI* or *PstI* and probed with the *GUS* gene. All lines for which both enzymes generated a single band were propagated to the T_3_ or T_4_ generation, and homozygous plants were used for quantitative comparisons. Expression levels were compared to the analogous intronless control pAR281 (37).

### Mapping transcription start sites with 5’-RACE

The 5’ ends of *TRP1:GUS* mRNAs were mapped by 5’-RACE (as described (40). Briefly, RNA extracted from transgenic plants was reverse transcribed (Reverse Transcription System; Promega, Madison, WI, USA) using the GUS-specific primer OAR37 (5’-TAACGCGCTTTCCCACCAACG-3’). The resulting cDNA was polyadenylated with terminal deoxynucleotidyltransferase and used as the template for PCR with the primers OAR37, QT (5’-CCAGTGAGCAGAGTGACGAGGACTCGAGCTCAAGCTTTTTTTTTTTTTTTTT-3’) and QO (5’-CCAGTGAGCAGAGTGACG-3’). A dilution of the first round PCR product was used as the template for a second round of PCR with the nested primers QI (5’-GAGGACTCGAGCTCAAGC-3’) and OAR29 (5’-GGTTGGGGTTTCTACAGGACG-3’). The 2nd round PCR product was cleaned up with a PCR Purification Kit (Qiagen, Hilden, Germany), digested with restriction enzymes *XhoI* and *BamHI*, and ligated into the vector pBluescript KS+. The DNA from transformants was screened by digestion with *SacI.* Clones with inserts long enough to contain the *SacI* site that begins 24 nt downstream of the *TRP1* start codon

## Acknowledgements

This research was supported by funding from the UC Davis PI Bridge Program. Jenna E Gallegos was supported by an NSF Graduate Research Fellows Program Grant 1148897.

